# Iatrogenic Iron Promotes Neurodegeneration and Activates Self-protection of Neural Cells against Exogenous Iron Attacks

**DOI:** 10.1101/2020.03.21.001925

**Authors:** Maosheng Xia, Shanshan Liang, Shuai Li, Zexiong Li, Manman Zhang, Beina Chen, Chengyi Dong, Binjie Chen, Ming Ji, Wenliang Gong, Dawei Guan, Alexei Verkhratsky, Baoman Li

## Abstract

Metal implants are used worldwide, with millions of metal nails, plates and fixtures grafted during orthopaedic surgeries. Iron is the most common element of these metal implants. As time passes metal elements can be corroded and iron can be released from the implants in the form of ferric (Fe^3+^) or ferrous (Fe^2+^). These iron ions can permeate the surrounding tissues and enter circulation; importantly both Fe^3+^ and Fe^2+^ freely pass blood brain barrier (BBB). Can iron from implants represent a risk factor for neurological diseases? This remains an unanswered question. In this study, we discovered that the probability of metal implants delivered through orthopaedic surgeries was higher in patients of Parkinson’s diseases (PD) or ischemic stroke than in healthy subjects. This finding instigated subsequent study of iron effects on neuronal cells. In experiments *in vivo*, we found that iron selectively decreased presence of divalent metal transporter 1 (DMT1) in neurones through increasing the expression of Ndfip1, which degrades DMT1 and rarely exists in glial cells. At the same time iron accumulation increased expression of DMT1 in astrocytes and microglial cells and triggered reactive astrogliosis and microglial activation. Facing the attack of excess iron, glial cells act as neuroprotectors to uptake more extracellular iron by up-regulating DMT1, whereas neurones limit iron uptake through decreasing DMT1 operation. Cerebral accumulation of iron was associated with impaired cognition, locomotion and mood. Excess iron thus affects neural cells and could increase the risk of neurodegeneration.

## INTRODUCTION

Every year, millions of metal implants such as metal nails, metal plates and other fixtures are used for orthopaedic intervention. At present, the follow-up mainly focuses on the operative prognosis, functional recovery, complications and secondary pathologies. However, effects of metal implants on nervous system and neurological conditions diseases are rarely considered. Metal implants are mainly composed from pure iron or iron-based alloys, the latter being more widely adopted in clinical practice. The common metal alloys contain Mn, Ti, Mg, Al, Co, Si etc., yet iron remains the primary element. Iron-based implants in the body they can be corroded and iron in the form of ferric or ferrous (Fe^3+^ or Fe^2+^) may permeate in the surrounding tissues and enter circulation; of note Fe^3+^ and Fe^2+^ can be mutually conversed by ferroxidase or ferrireductase.^1–3^ Corrosion of metallic components in the tissues may trigger inflammation and accumulation of iron in macrophages in months or years following the implantation.^1,2^ Manipulations with iron content in rodents through the diet, intraperitoneal (i.p.) injection or subcutaneous infusion lead to iron deposition in different tissues and in plasma.^4^

Ionised iron crosses blood-brain barrier and blood-cerebrospinal barrier.^5^ Aberrant iron accumulation in the brain has been reported to be associated with neurodegenerative diseases, such as Alzheimer’s diseases (AD), Parkinson’s diseases (PD), and Huntington diseases.^6–8^ Oxidative stress resulting from the accumulated iron triggers neuronal death leading to neurodegeneration;^9^ iron being associated with production of reactive oxygen species (ROS), which injure cellular membranes, proteins and deoxyribonucleic acid (DNA).^10^ Divalent metal transporter 1 (DMT1/SLC11A2) is an H^+^-driven multi-metal transporter responsible for transmembrane iron transport;^11^ in particular DMT1 has a principle role in transporting iron into the brain.^12^ In the central nervous system (CNS), DMT1 is widely expressed in neurones, astrocytes, microglial cells, endothelial cells and oligodendrocyte progenitors.^13^ How iron affects all these cells *in vivo* remains mostly unknown.

In this study, we performed retrospective analyses of patients with the history of metal implants; in particular we focused on patients diagnosed with PD, ischemic-stroke or neurologic tumours (a disease-related control group). We discovered that metal implantation increased the risk of occurrence of PD and ischemic-stroke. In the animal model, we studied how exogenous iron acts on neural cells, and we found the reverse effects of iron on the levels of DMT1 in neurones and glia. Glial cells respond to exogenous iron attack by up-regulating DMT1 expression, whereas neurones accelerated DMT1 degradation; both these processes being arguably neuroprotective. Our study suggests that iron intake (including that associated with orthopaedic implants) should be considered as a risk factor for neurological disorders.

## RESULTS

### High incidence of metal implantation in PD and ischemic stroke

Table 1 shows the general characteristics of the subjects under study. There was no significant difference in age-distribution between control group and any of disease groups (Supplementary Fig. 1A). Similarly, there was no gender-distribution difference, and the male-female ratio was close to 1.0 in all groups. There was no difference in educational level, ethnical origin (individuals of Han race accounted for more than 96.43% in control or any case group, which was matched to the national population percentage of Han race in China). The incidence of number of metal inserts was 3.60%, *X*^2^(1)=8.307, p=0.004 in PD group, 5.00% (*X*^2^(1)=16.225, p<0.0001) in ischemic-stroke group, while it was only 1.14% in healthy control group. In the cerebral tumour group the insert incidence was 1.50% (p=0.716) which was not significantly different from healthy control group.

**Table 1.**
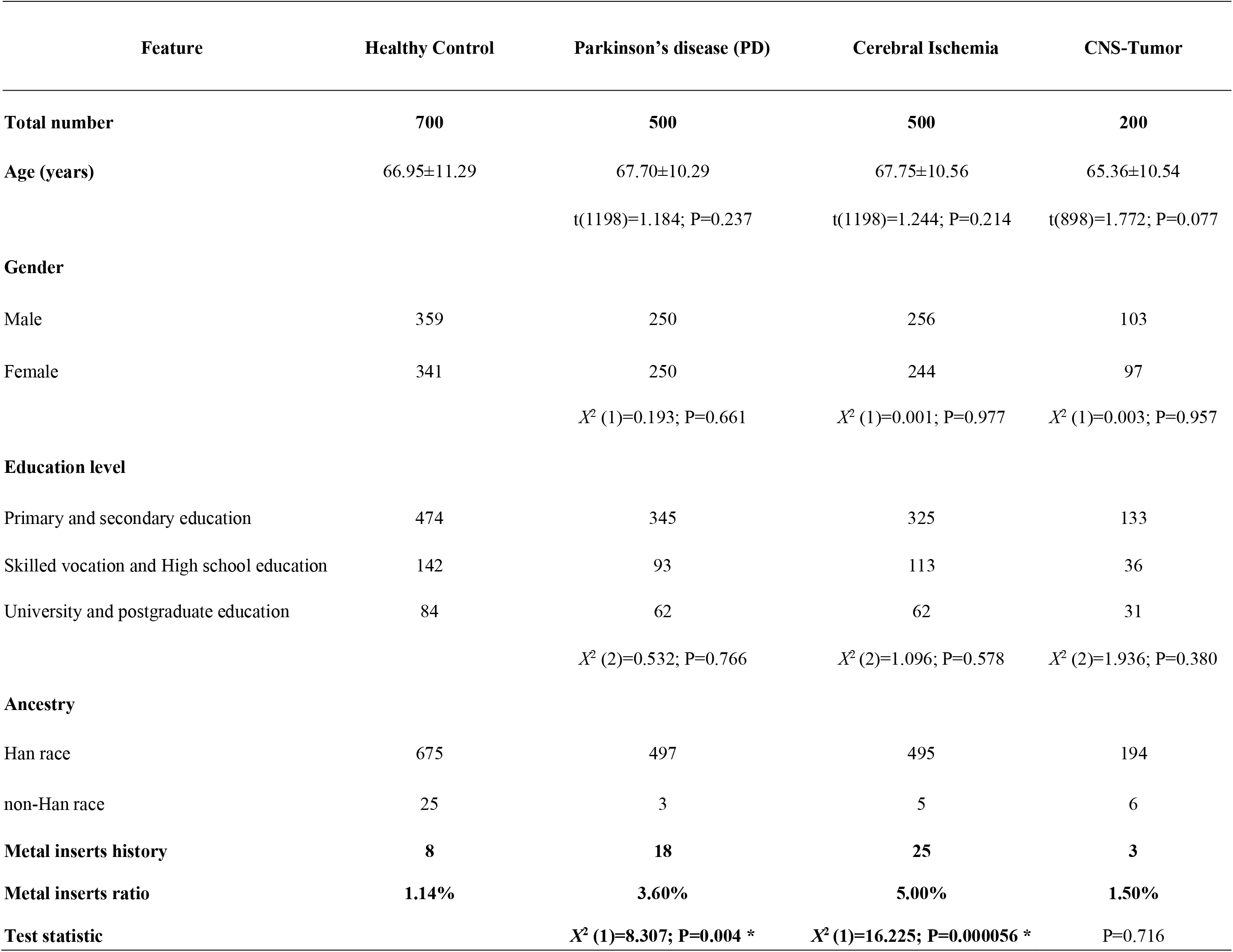
General characteristics of 500 Parkinson’s disease cases, 500 cerebral ischemia cases, 200 CNS-Tumor cases and 700 healthy controls

Detailed distribution of cases having inserts were further analysed; with the results shown in Supplementary Table 1. In control group, 8 out of 700 individuals had metal implants history; in PD group 18 subjects out of 500 had metal inserts, whereas in ischemia group 25 patients out of 500 had metal implants. According to the age at the moment of diagnosis, all subjects were divided into four age groups, <46, 46-60, 61-75 and >75 years. Comparing to the age distribution in control group, the metal inserts cases were more frequent in the last two groups (Supplementary Fig. 1B). In PD group, 11 out of 244 patients from 61 - 75 years old group had inserts (*X*^2^(1)=4.414, P=0.036), in patients older than 75 years, 3 out of 38 underwent metal implantation 3/38 in >75 years old group (P=0.043). In ischemia group, the inserts frequencies were significantly different from healthy subjects in 3 age groups: 5 out of 111 in 46 – 60 years old group (P=0.024), 12 out of 259 in 61-75 years old group (P=0.029) and 8 out of 111 in patients older than 75 years (P=0.016). The distribution of gender, ethnicity and education showed no difference between all groups. Patients were also divided into three groups, according to the time between implant surgery to PD or ischemia diagnosis, <5, 5-10 and >10 years. The first group showed more cases with metal inserts diagnosed with PD or ischemia (Supplementary Fig. 1C), the incidences were 12 out of 18 (66.67%, P=0.002) and 13 out of 25 (52.00%, P=0.044), respectively.

### The high PD-occurrence after orthopaedic surgeries with metal implants

In Table 2, we present the occurrence of PD after orthopaedic surgeries with and without metal implants in the past ten years. Since 2009, we counted 7500 subjects who underwent orthopaedic surgeries without using metal implants and 15000 subjects which had orthopaedic surgeries with metal implantation. The occurrence of PD in surgeries without metal implants was 1.31%, whereas this ratio increased to 1.98%, in surgeries with metal implants; this difference reached the level of statistical significance (*X*^2^(1)=13.143, p<0.001).

**Table 2.**
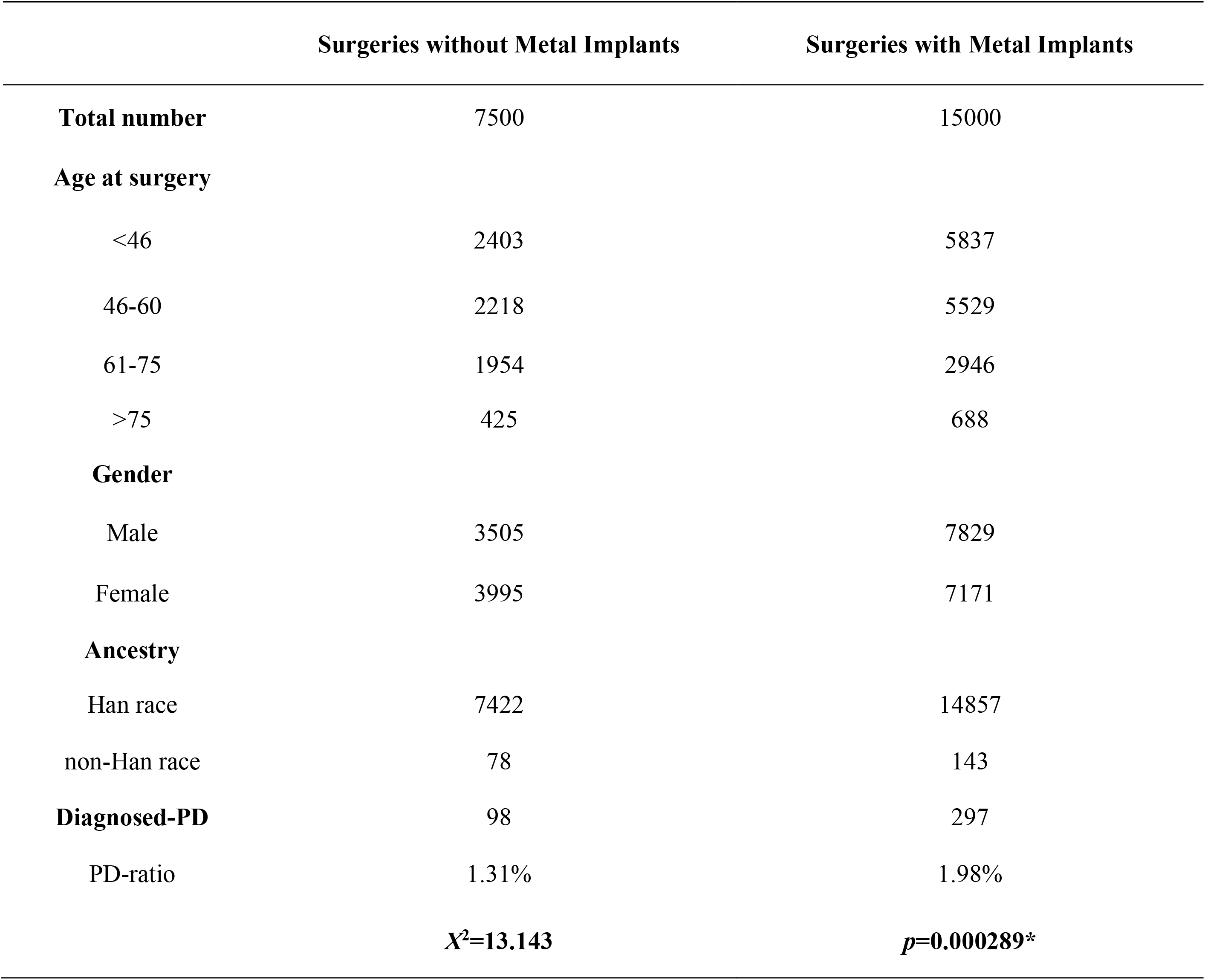
The occurrence of diagnosed PD in the orthopedic surgeries without and with metal implants.

|  | Surgeries without Metal Implants | Surgeries with Metal Implants |
| --- | --- | --- |
| **Total number** | 7500 | 15000 |
| **Age at surgery** |  |  |
| <46 | 2403 | 5837 |
| 46-60 | 2218 | 5529 |
| 61-75 | 1954 | 2946 |
| >75 | 425 | 688 |
| **Gender** |  |  |
| Male | 3505 | 7829 |
| Female | 3995 | 7171 |
| **Ancestry** |  |  |
| Han race | 7422 | 14857 |
| non-Han race | 78 | 143 |
| **Diagnosed-PD** | 98 | 297 |
| PD-ratio | 1.31% | 1.98% |
|  | **$X^2=13.143$** | **$p=0.000289^*$** |

The number of PD-diagnosed subjects and the total number of orthopaedic surgeries with or without metal inserts on yearly basis is shown in Fig. 1. During the period between 2009 and 2011, we found significant difference between using metal implants surgeries group (inserts group) and orthopaedic surgeries without implants (control group). In 2009, 7 subjects were diagnosed with PD out of 567 subjects having orthopaedic surgeries, but the PD occurrence was 45 out of 1646 in metal inserts group (*X*^2^(1)=4.132, P=0.042). In 2010, the PD occurrences in control and inserts groups were 11 out of 530 and 37 out of 893 respectively (*X*^2^(1)=4.363, P=0.037). In 2011, the PD occurrence in these two groups was 10 out of 656 and 43 out of 1418 (*X*^2^(1)=4.096, P=0.043). In the ensuing seven years, there was no significant difference in the occurrence of PD between control and inserts groups.

**Figure 1.**
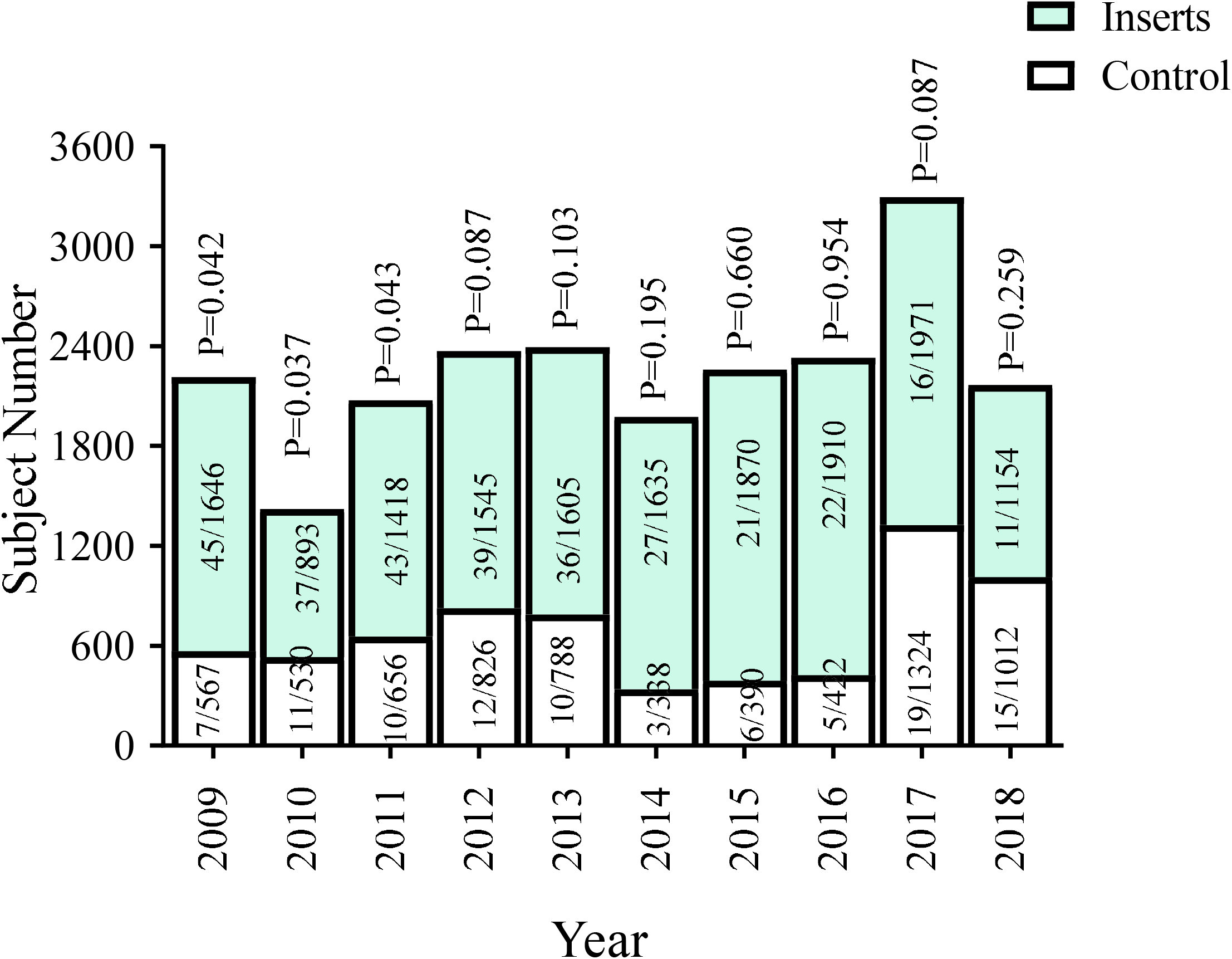
The occurrence of PD after taking orthopaedic surgeries from 2009 to 2018. The numbers of the subjects and PD diagnoses after orthopaedic surgeries with and without using metal implants.

### Iron overload reversely regulates expression of DMT1 in neurones and glia

To study the effects of iron in animals we injected mice hypodermically with iron dextran. Iron accumulation was identified in the brain using Perl’s staining. Iron accumulation increased with an increase in the intravenously injected dose (Fig. 2A). Four cerebral regions showed iron accumulation following 2 mg iron injections in FC, HP, SN and ST (Fig. 2B). The expression of DMT1 in astrocytes around big vessels was also detected in FC and SN (Fig. 2C-2E). Iron increased the immunofluorescence intensity of DMT1 to 224.42 ± 11.49% (p < 0.001, n = 6) of control group in FC (Fig. 2E), and to 215.42 ± 7.20% (p < 0.001, n = 6) of control group in SN (Fig. 2E).

**Figure 2.**
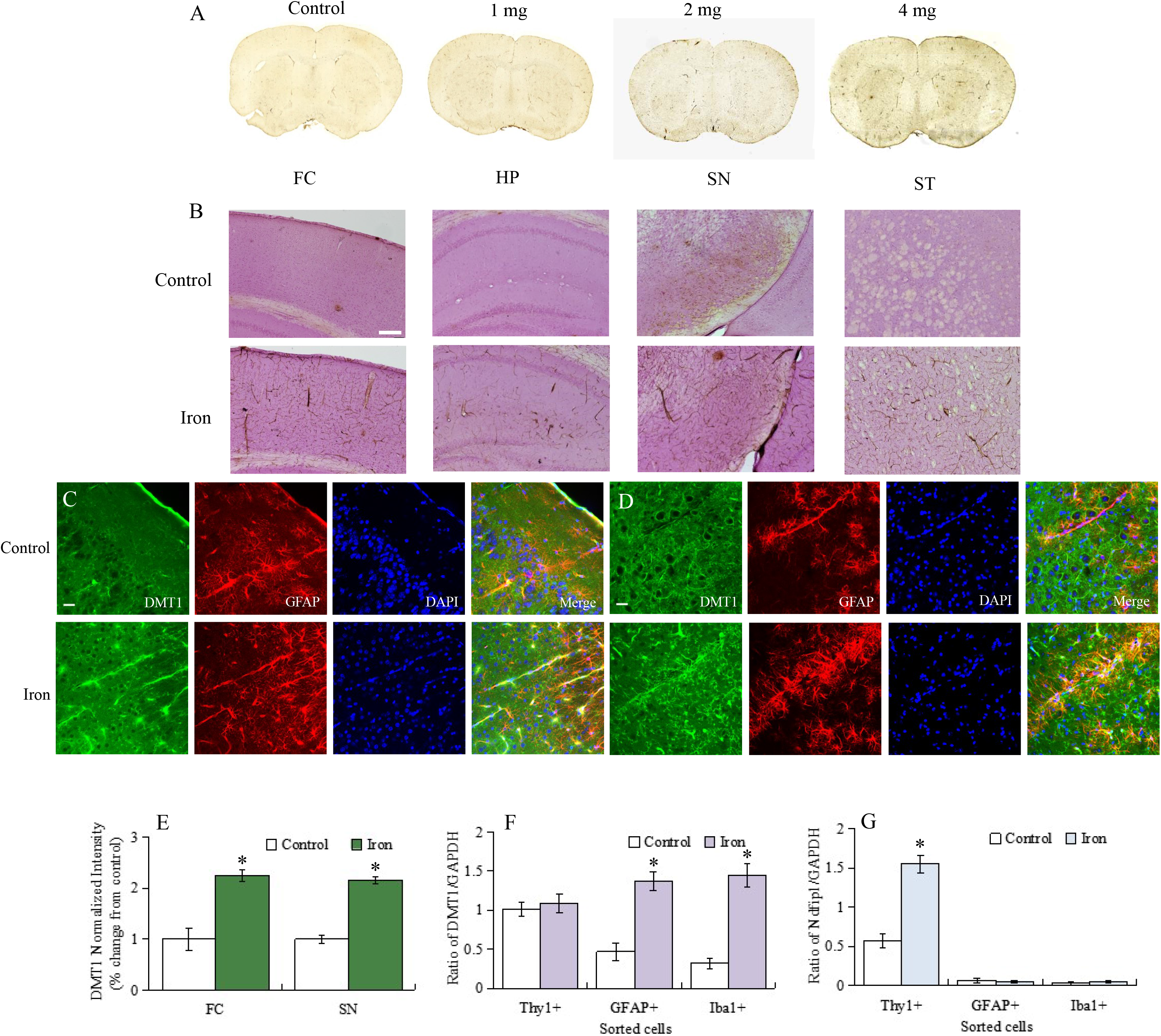
Iron accumulation in the brain and expression of DMT1 specific mRNA. (**A**) Perl’s images of the whole brain slice treated with different concentrations of iron dextran (0, 1, 2 and 4 mg/kg/day) for 6 days. (**B**) Perl’s images of frontal cortex (FC), hippocampus (HP), substantia nigra (SN) and striatum (ST) with the absence or presence of 2 mg/kg/day iron dextran. Scale bar, 500 μm. (**C and D**) Immunofluorescence images of DMT1 distributed in astrocytes around vessels after treatment with 2 mg/kg/day iron dextran. DMT1 (green), GFAP (red), DAPI (blue) and merged images were respectively shown in FC (**C**) and SN (**D**). (**E**) The fluorescence intensity of DMT1 co-stained with GFAP represent mean ± SEM, n = 6. Scale bar, 25 μm. *p < 0.05, statistically significant difference from control group. (**F and G**) The relative mRNA expression ratio of DMT1 and Ndfip1. After treatment with 2 mg/kg/day iron dextran for 6 days, neurones (Thy1+), astrocytes (GFAP+) or microglial cells (Iba1+) were FACS sorted and selected for qPCR measurement. The relative mRNA expression ratio of DMT1/GAPDH (**F**) and Ndfip1/GAPDH (**G**) represent mean ± SEM, n=6. *p < 0.05 - statistically significant difference from control group.

Expression of mRNA for DMT1 and Ndfip1 was determined in neurones, astrocytes and microglial cells (Fig. 2F, G). In FACS-sorted neurones, treatment with iron dextran did not significantly change mRNA expression of DMT1, but increased the mRNA level of Ndfip1 to 271.93 ± 11.12% (p < 0.001; n = 6) of control group. Administration of iron dextran up-regulated mRNA expression of DMT1 to 291.49 ± 12.37% (p < 0.001; n = 6) and 453.13 ± 14.97% (p < 0.001; n = 6) of control group in astrocytes and microglial cells, respectively. However, iron treatment did not change mRNA expression of Ndfip1 in glial cells.

Treatment with iron dextran decreased fluorescent intensity of DMT1 immunostaining in neurones but increased it in astrocytes and microglia (Fig. 3A-O). In FC, for example, when compared with control group, the fluorescence intensity of DMT1 in neurones decreased to 47.54 ± 4.88% (p < 0.001, n = 6), whereas it increased to 143.92 ± 10.19% (p = 0.021, n = 6) and to 195.30 ± 8.84% (p < 0.001, n =6) in astrocytes and microglia (Fig. 3M-3O). Similar intensity changes of DMT1 were also observed in other three cerebral regions. The treatment with iron dextran decreased the fluorescence intensity of DMT1 in neurones to 22.93 ± 5.04% (p = 0.009, n = 6), 54.74 ± 3.90% (p < 0.001, n = 6) and 46.24 ± 6.48% (p = 0.006, n = 6) in HP, SN and ST, respectively. In these three brain regions, the intensity of DMT1 was increased to 162.57 ± 11.11% (p < 0.001, n = 6), 169.54 ± 5.57% (p < 0.001, n = 6) and 189.94 ± 21.97% (p = 0.001, n = 6) in astrocytes, and it was elevated to 223.58 ± 12.15% (p < 0.001, n = 6), 142.82 ± 6.01% (p < 0.001, n = 6) and 127.62 ± 6.50% (p = 0.002, n = 6) in microglia, compared with control group (Fig. 3M-O).

**Figure 3.**
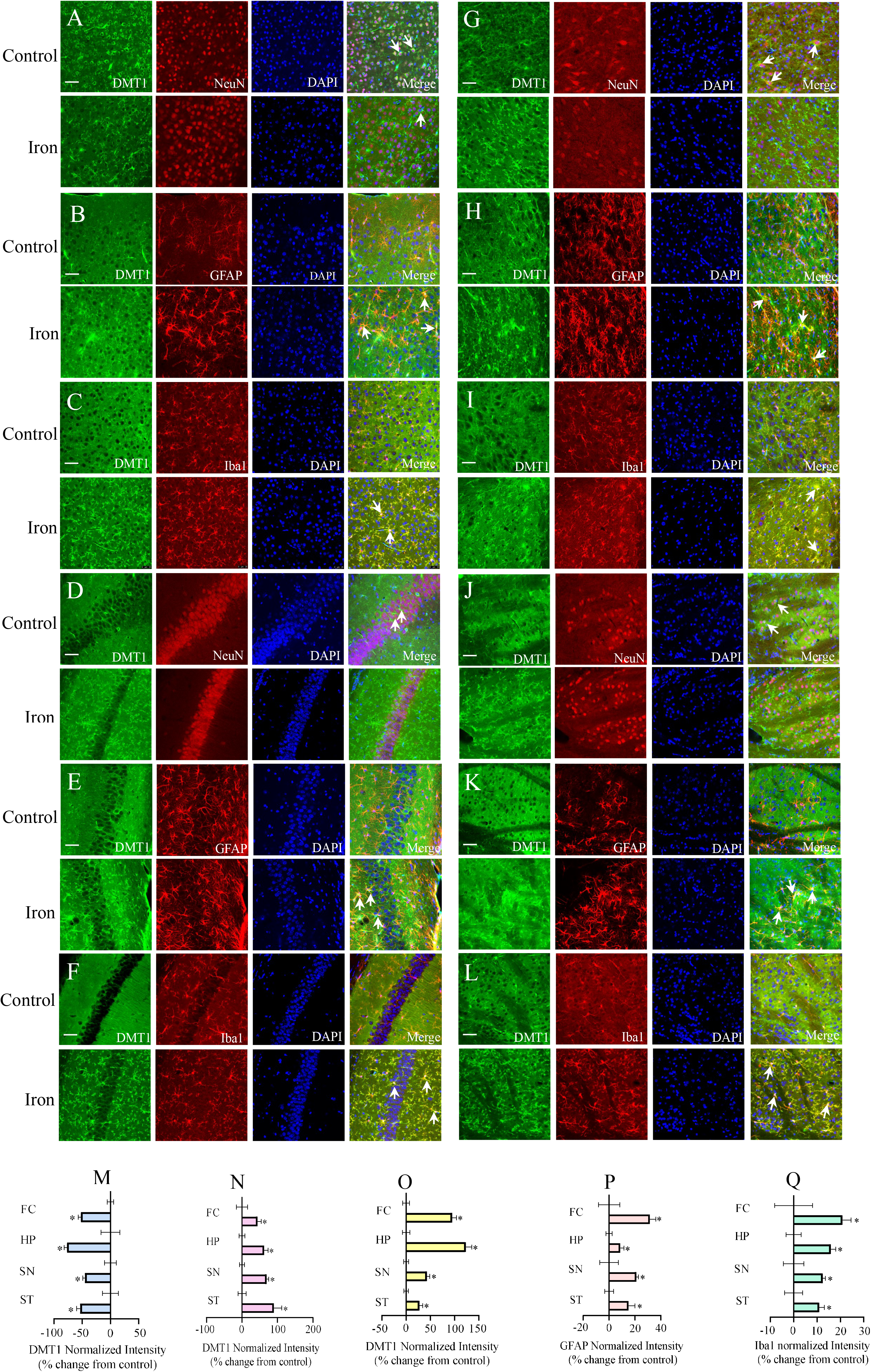
Protein levels of DMT1 in neurones and glia in four cerebral regions. **(A-L)** After treatment with 2 mg/kg/day iron dextran for 6 days, the images of DMT1 (green) co-stained with NeuN, GFAP or Iba1 (red) and DAPI (blue) were observed in FC (**A**-**C**), HP (**D**-**F**), SN (**G**-**I**) and ST (**J**-**L**). (**M**-**O**) The fluorescence intensity of DMT1 merged with NeuN, GFAP or Iba1 were separately normalised by control group and represent mean ± SEM, n = 6. (**P and Q**) The fluorescence intensity of GFAP and Iba1 were normalized by control group and represent mean ± SEM, n = 6. Scale bar, 50 μm. *p<0.05, statistically significant difference from control group.

### Iron accumulation triggers reactive gliosis

Iron dextran significantly increased the fluorescence intensity of GFAP and Iba1 in four cerebral regions (Fig. 3P and 3Q). Compared with control group, the intensity of GFAP was significantly increased to 131.48 ± 4.59% (p = 0.008, n = 6), 108.71 ± 2.79% (p = 0.040, n = 6), 121.16 ± 1.50% (p = 0.017, n = 6), 115.03 ± 4.94% (p = 0.031, n = 6) (Fig. 3P), the intensity of Iba1 was elevated to 120.91 ± 3.68% (p = 0.041, n = 6), 115.96 ± 2.06% (p = 0.002, n = 6), 112.42 ± 1.18% (p = 0.022, n = 6), 110.93 ± 2.23% (p = 0.035, n = 6) in FC, HP, SN and ST, respectively (Fig. 3Q). Iron dextran changed the morphology of astrocytes labelled with GFAP indicative of reactive astrogliosis (Supplementary Fig. 2). Meanwhile, ferrous sulfate (FeSO_4_) at 1 mM triggered intracellular calcium increase in primary cultured astrocytes (Supplementary Fig. 3).

### Iron accumulation instigates neuronal apoptosis and abnormal behaviours

The ratio of TUNEL+/DAPI+ (total cells) was elevated to 243.45 ± 21.06% (p < 0.001, n = 6), 241.94 ± 26.82% (p < 0.001, n = 6) and 237.75 ± 19.59% (p < 0.001, n = 6) of control group in FC, HP and SN; there was no significant difference between control and iron dextran groups in ST region (Fig. 4A-E). Treatment with iron dextran significantly increased the level of ROS to 190.69 ± 28.41% (p = 0.017, n = 6) in FC, to 171.39 ± 15.75% (p = 0.026, n = 6) in HP, to 164.01 ± 16.42% (p = 0.031, n = 6) in SN, as comparison with control group (Fig. 4F). However, in ST, iron dextran did not affect the level of ROS.

**Figure 4.**
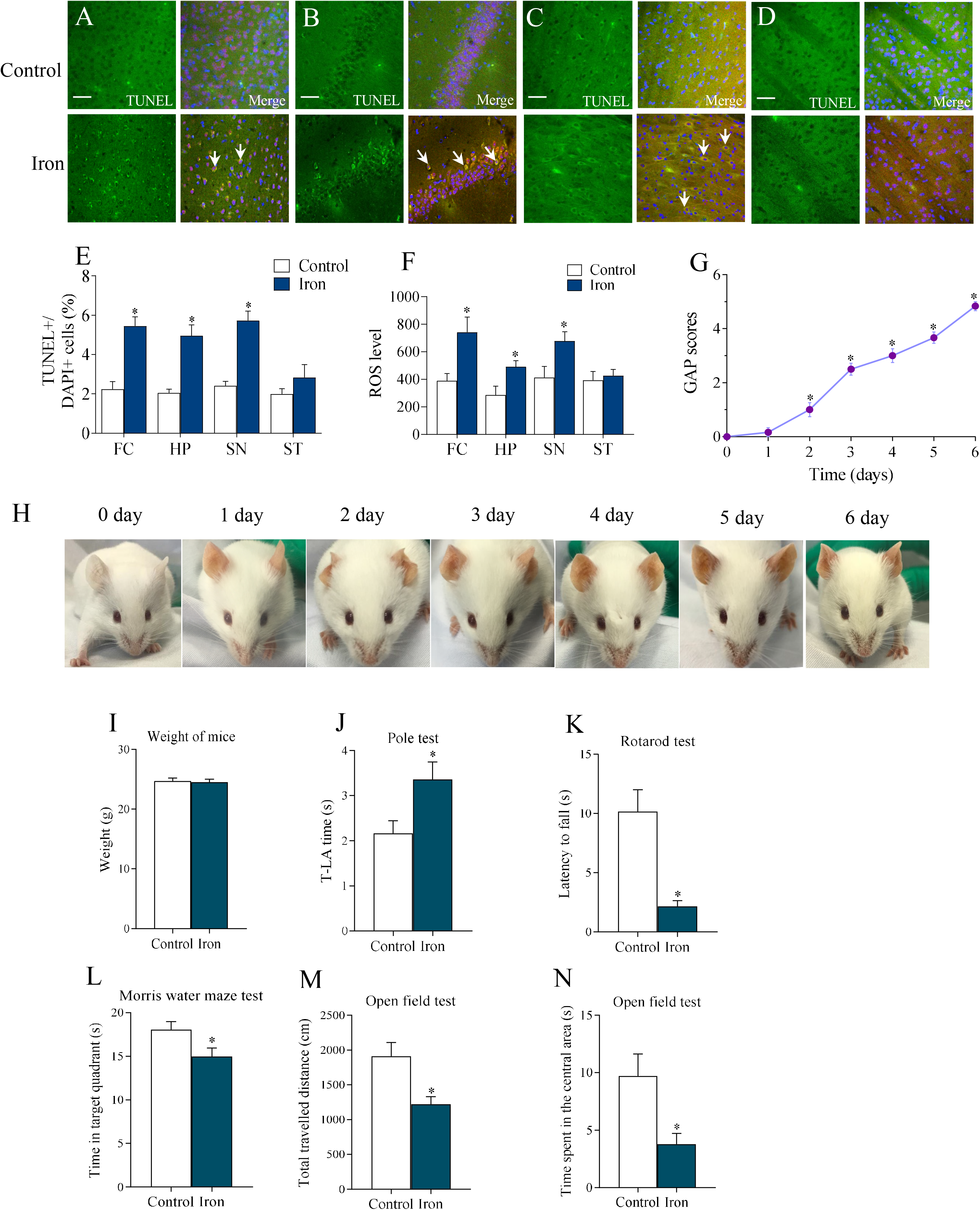
Effects of excess iron on animals behavior. **(A-D)** After treatment with 2 mg/kg/day iron dextran for 6 days, TUNEL assays were measured in FC (**A**), HP (**B**), SN (**C**) and ST (**D**). Meanwhile, the neurons were stained with NeuN (red) and nuclei were labelled with DAPI (blue). (**E**) Ratios of TUNEL+/DAPI+ cells presented as mean ± SEM, n = 6. Scale bar, 50 μm. *p < 0.05, statistically significant difference from control group. (**F**) ROS level in these four cerebral regions. Data represent mean ± SEM, n = 6. *p<0.05, statistically significant difference from control group. (**G**) GAP scores treatment with 2 mg/kg/day iron dextran from 0 to 6 days. Data represent mean ± SEM, n=6. *p<0.05, statistically significant difference from 0 day. (**H**) The appearances of mice treatment with 2 mg/kg/day iron dextran from 0 to 6 days. (**I**) Changes in mice weight. (**J**) The movement time from the pole top to the floor (T-LA time) was measured in pole test. (**K**) The dwelling time on rotating bar was recorded in rotarod test. (**L**) Time spent in target quadrant was measured in Morris water maze test. (**M and N**) The total travelled distance (**M**) and time spent in the central area (**N**) were analyzed in open field test. Data represent mean ± SEM, n=6. *p<0.05, statistically significant difference from control group.

The general appearances of mice are shown in Fig. 4G, H. The score of mice in absence of iron was 0, the GAP scores were gradually increased during 6 days treatment with iron dextran, the score was increased to 4.83 ± 0.17 (n = 6) at day 6. During the administration of iron dextran, the mice gradually showed a series of abnormal behaviours, such as restlessness, guarding, arched back, irregular breath and segregation. Specially, the eyes were glazed and sunken, and the colour of ears darkened (Fig. 4H).

Meanwhile, the relevant behavioural tests were also performed. The average weight of mice was not changed by iron injection (Fig. 4I). In pole test, iron dextran prolonged the used T-LA time to 154.95 ± 17.96% of control group (p = 0.035, n = 6) (Fig. 4J). The dwell time in rotarod test was shorten by iron dextran to 21.31±4.69% of control group (p = 0.002, n = 6) (Fig. 4K). Administration of iron dextran significantly decreased the time spent in target quadrant in Morris water maze test to 83.00 ± 5.33% of control group (p = 0.029, n = 6) (Fig. 4L and Supplementary Fig. 5). In open field test, the travel distance and time spent in central area were decreased to 63.98 ± 5.65% (p = 0.010, n = 6) and 38.97 ± 9.78% (p = 0.017, n = 6), compared with control group (Fig. 4M and 4N).

**Figure 5.**
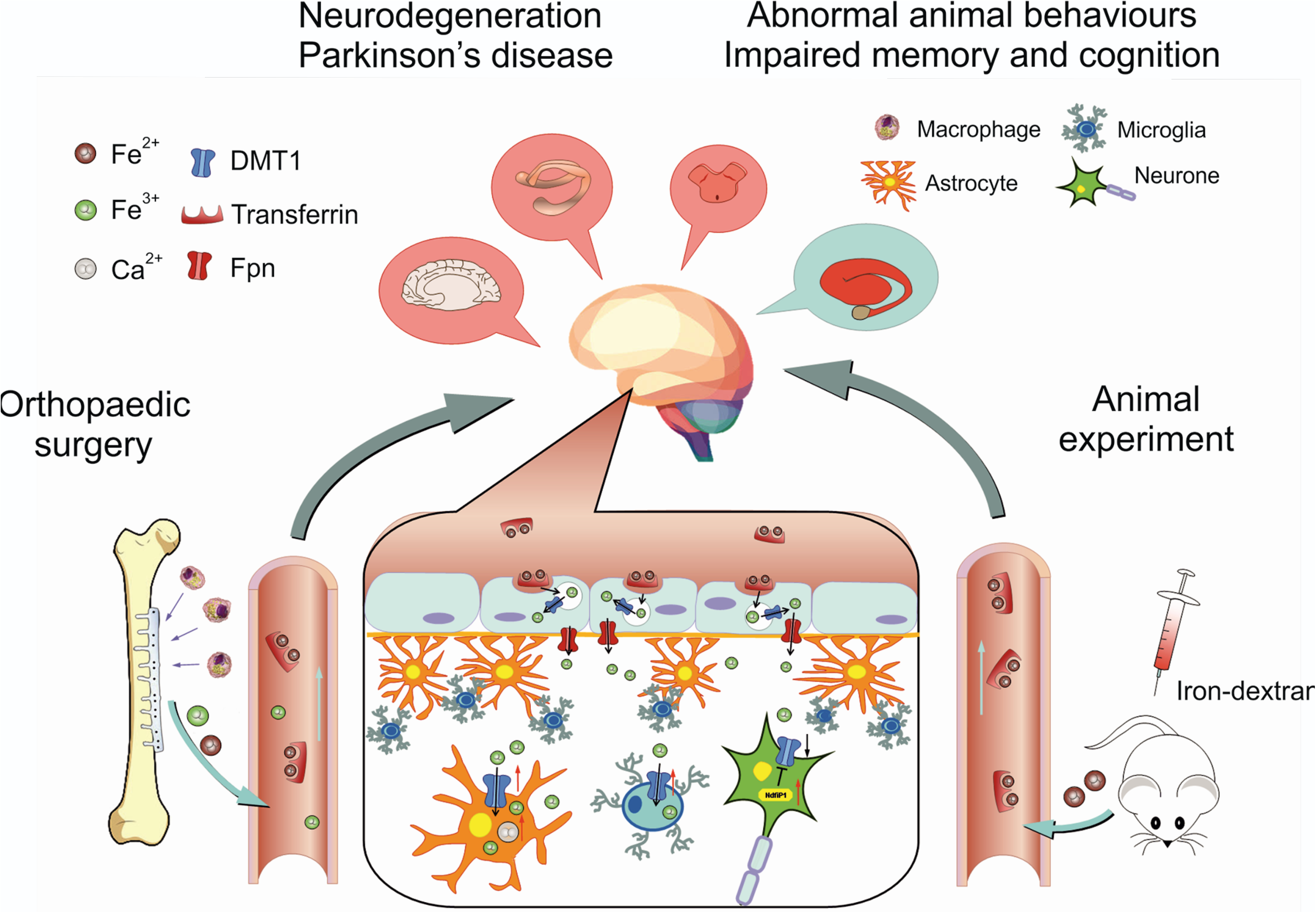
Adverse effects of excess iron on neural cells. After inserting metal implants in orthopaedic surgery, or after injections of iron-dextran to experimental animals the ionised Fe^2+^/Fe^3+^ enters the blood stream. Through transportation of transferrin and the ferroportin (Fpn), the Fe^2+^/Fe^3+^ crosses blood-brain barrier and blood-cerebrospinal barrier and accumulates in the brain. Facing the attack of excess iron, glial cells (astrocytes and microglia) act as protectors by increasing the uptake of ferrous by up-regulating DMT1, the main iron transporter. At the same time neurones restricts the influx of iron by decreasing expression of DMT1 by up-regulation of Ndfip1. This phenomenon has been observed in neurodegenerative cerebral regions, FC, HP, SN and ST. The increased apoptosis of neurones and reactive gliosis may instigate abnormal animal behaviours as well as impair memory and cognition.

## DISCUSSION

Iron is a necessary element in the human body, the iron deficiency is harmful for development and health, but the excessive iron can also impair subjects’ health. According to our study, and as summarised in Fig. 5, the application of iron-based metal implants in the orthopaedic surgeries increased the occurrence risk of PD and ischemic stroke. Specifically, implantation surgeries increased probability of PD or ischemia in the five years period after intervention. However, in this study, we focused on the effects of excessive iron on impairments of neuronal cells in PD context. We will further study the effects of iron on the high risk of ischemia in future research, focusing on possible action of iron on vascular endothelial cells. The confirmed age for patients diagnosed with PD and having metal implants tended to be more than 60 years old, which conformed with the peak of onset age in this disease. The occurrence risk of PD was significantly increased in the metal implants group compared with the absence of metal implants group that have other orthopaedic surgeries.

To analyse underlying mechanisms of iron effects on CNS, we studied effects of hypodermic injections of iron dextran on the expression of DMT1. We discovered that treatment with iron changed protein expression of DMT1 in neurones and glial cells in an opposite manner. In FC, HP, SN and ST, neuronal DMT1 was decreased by iron, whereas iron significantly enhanced expressions of DMT1 in astrocytes and microglial cells. We speculated that this phenomenon reflects a neuroprotective mechanism activated by iron attack. As shown in Fig. 5, for resisting the invasion of iron, glial cells act as protectors, as they increase the uptake of ferrous by up-regulating DMT1, in particular in astrocytes distributed along the big vessels. At the same time neurones restrict the influx of iron by decreasing DMT1 levels. Decreased protein level of DMT1 induced by iron in neurones was not due to the downregulation of mRNA, but resulted from increased degradation by Ndfip1, an adaptor protein for Nedd4 family of ubiquitin ligases,^14,15^ because the mRNA of Ndfip1 was significantly and selectively increased by iron in neurones. The chronic treatment with iron can increase the expression of Ndfip 1 in MES23.5 cells,^16^ but Ndfip1 is not expressed in glial cells.^17^ Our results also support that Ndfip1 rarely existed in glial cells with or without iron treatment.

Chronic treatment with iron triggered ROS production in the cerebral regions of FC, HP and SN, but not in ST. Neuronal apoptosis induced by iron and mediated by ROC was also identified in these three regions. Behavioural tests demonstrated that treatment with iron evoked anxiety behaviours and resulted in cognitive and movement dysfunction. The extent of oxidative stress triggered by iron was different in dopaminergic and cholinergic neurones that was reflected by the difference of apoptosis level between SN and ST. Early and prominent neuronal apoptosis observed in SN compared to the striatum may explain the clinical symptoms of PD associated with relatively hyperactive acetylcholine function. Following treatment with iron dextran cerebral deposition of iron was gradually increased and the mice developed abnormal appearance changes, poor coordination and locomotion, declined cognition and increased anxiety-like behaviours. These changes are somewhat resembling the higher dementia rate after fractures in clinic.^18^

In addition, reactive gliosis (as indicated by an increase in expression of GFAP and Iba1) was induced by iron. We also discovered that exposure to ferrous triggers transient increase in intracellular Ca^2+^, and this Ca^2+^ signals may be a trigger of astrogliosis. Hence, facing the invasion of excess iron, glial cells undergo activation and protect neurones by taking up excess of extracellular iron.

In conclusion, neuronal apoptosis and reactive gliosis both accompanied iron overload which played a role of pathology promoter. The excessive iron in bodies should be of special concern and the patients taking iron-based metal implants in orthopaedic surgeries should be considered at risk for the neurodegenerative diseases, because excess iron may injure neural tissue.

## METHODS and MATERIALS

### Populational analysis

We firstly set up a case-control study with common cerebral diseases, including 500 patients diagnosed with PD, 500 cases of ischemic stroke, 200 cases of brain tumour (glioma, meningioma and hypophysoma) and 700 healthy controls. Cases having tumour were used as the disease-related control group. Then, for investigating the effects of orthopaedic surgeries with and without using metal implants on the risk of PD, we set up a case-cohort study with the inpatients from orthopaedic departments, including 15,000 patients from metal implant surgeries and 7,500 patients from solo orthopaedic surgeries. The anamneses were collected from the First Affiliated Hospital of China Medical University, which is the biggest general public hospital in the northeast of China and has about 3 million outpatients and 159,000 inpatients every year. We selected cases of inpatients with definitive diagnosis from 2009 to 2018. Population sources were from the four provinces of China (Heilongjiang, Jilin, Liaoning and Neimenggu). The individuals in healthy control group were similarly selected from these four provinces. This study was authorised and approved by Medical Ethics Committee of China Medical University, No. [2019]059.

### Data collection

The collected data included age, gender, race, educational background, contact address and the history of iron-based alloys implants. Among these factors, age and the time from metal implant surgery to be diagnosed related disease were quantified. The inclusion criteria were (i) ages more than 18-year old, (ii) PD diagnosis-patients were diagnosed according to UK PD Society Brain Bank Clinical Diagnostic,^19^ (iii) ischemic stroke - diagnosed by CT or MRI images, (iv) brain tumour - identified by histological analysis of tissues removed from neurosurgery excision. Healthy controls were selected from age-matched individuals to case-group, who did not have neurodegenerative diseases, stroke, brain tumour and other cerebral diseases. For all groupings, the exclusion criteria were: (i) the patients had any psychiatric diseases or had two or more cerebral diseases together; (ii) the patients had been diagnosed cerebral diseases before the orthopaedic surgeries; (iii) the patients had the history of any other surgery except this study-relevant orthopaedic surgeries or neurosurgeries; (iv) the patients lost contact or had the difficulty of language communication.

### Animals

Male wild type mice, B6.Cg-Tg(Thy1-YFP)HJrs/J and FVB/N-Tg(GFAP-GFP)14Mes/J mice were obtained from Jackson Laboratory (Bar Harbor, ME, USA. Mice were 10 - 12 weeks old (~25 g) and were kept in standard housing conditions (22±1°C; light/dark cycle of 12 h/12 h) with food and water available ad libitum. All operations were carried out in accordance with the USA National Institutes of Health Guide for the Care and Use of Laboratory Animals (NIH Publication No. 8023) and its 1978 revision, and all experimental protocols were preregistered and approved by the Institutional Animal Care and Use Committee of China Medical University.

### Materials

Most chemicals, including iron dextran (ferric hydroxide dextran complex) and 4’, 6-diamidine-2’-phenylinedole dihydrochloride (DAPI) were purchased from Sigma (MO, USA). The primary antibody raised against DMT1 was purchased from Santa Cruz Biotechnology (CA, USA). Primary antibody raised against NeuN, GFAP and secondary antibodies was purchased from Thermo Fisher Scientific (CA, USA). Primary antibody raised against Iba-1 was purchased from Wako Pure Chemical Industries (Osaka, Japan). Micro serum iron concentration assay kit, potassium ferrocyanide and diaminobenzidine (DAB) kit were purchased from Solarbio Life Sciences (Beijing, China). TUNEL apoptosis detection kit was purchased from Roche-Sigma (USA). The ROS assay kit was purchased from BestBio (Shanghai, China).

### Iron-treatments

Iron dextran was hypodermically injected at 200 μl volume containing 1 mg, 2 mg or 4 mg/kg/day for 6 days, dextran free from iron was injected the same volume in control group.

### Immunohistochemistry

Brain tissue was immersed in 4% paraformaldehyde and cut into 50 μm slices. As our previous described,^20,21^ the slices were pre-incubated with normal donkey serum (NDS, 1:20; Jackson Immuno-Research Laboratory) for 1 h and a mixture of primary antibodies overnight, mouse anti-DMT1 (1:100), rabbit anti-NeuN (1:200), rabbit anti-GFAP (1:200) or rabbit anti-Iba1 (1:200). After several rinses, the slices were incubated with Alexa fluor-conjugated secondary antibodies for 2 hours at room temperature, DAPI (1:1000) was stained to identify cell nuclei. Immunofluorescence was imaged using a confocal scanning microscope (DMi8, Leica, Germany). To measure the level of DMT1 in neurones, astrocytes and microglia, we calculated the mean intensity of DMT1 immunofluorescence which was co-labelled with NeuN, GFAP or Iba1 in cortical, hippocampus, substantia nigra and striatum regions. The background intensity of each image was collected in cell-free parenchyma in the same field of view and subtracted from the immunofluorescence intensity. The intensity of DMT1 immunofluorescence from each group was normalised to the intensity of the control group.

### Perl’s staining

Tissue iron accumulation was identified by Perl’s staining with modifications. The slices were immersed in the mixed reaction liquid (4% potassium ferrocyanide: 4% hydrochloric acid = 1:1) at 37°C for 30 minute and then stained using a DAB kit for 4 min. The observations were imaged using an optical microscope (BX53, Olympus, Japan) or Carl Zeiss Promenade 10, Jena, Germany).

### Sorting neural cells through fluorescence activated cell sorter (FACS)

To measure the mRNA of DMT1 and Ndfip1, neurones expressed fluorescent marker YFP (Thy1-YFP mice), whereas astrocytes expressed GFP (GFAP-GFP mice). As previously described,^22,23^ the cells isolated from wild type mice were labelled by Iba1 positive antibody, and the cells from transgenic mice were used for sorting specific neurons or astrocytes populations by FACS. Dead cells were excluded based on positive DAPI signals. The cells were sorted in cold Minimum Essential Media (MEM) containing 1% Bovine Serum Albumin (BSA). The RNA of the sorted cells was extracted by Trizol.

### Quantitative PCR (qPCR)

Total RNA was reverse transcribed and PCR amplification was performed in a Robo-cycler thermocycler, as our previous description.^20,22^ Briefly, relative quantity of transcripts was assessed using five folds serial dilutions of RT product (200 ng). RNA quantity was normalised to glyceraldehyde 3-phosphate dehydrogenase (GAPDH) before calculating relative expression of DMT1 or Ndfip1. Values were expressed as the ratio of the relative expression of DMT1/GAPDH or Ndfip1/GAPDH.

### Calcium Imaging

Primary cultured astrocytes were incubated in cell-permeant Fluo4-AM (Invitrogen) for 30 min at 37°C and 30 min at room temperature. As described previously,^24,25^ dye-loaded cells were excited at 488 nm and emission was measured at wavelength of 522 nm. Images were acquired every 6 seconds. Calcium transients were initiated by the addition of 1 mM FeSO_4_ after 1 minute. Images were acquired using confocal scanning microscope (DMi8, Leica, Germany) and analyzed using ImageJ software (National Institutes of Health, Bethesda, MD). F_0_ (baseline) and F are the mean fluorescence intensities, the fluorescence intensity was calculated as the percentage of F/F_0_.

### Ethogram

As described previously,^26^ we assessed the mice normal and abnormal behaviours according to general appearance parameters (GAP) assessments. A score of 0 or 1 for the categories of activity, posture, breathing pattern, coat condition, and interaction with other mice were given.

### Morris water maze test

As our previous description,^20^ the Morris water maze test was carried to estimate a spatial learning and memory performance. The mice were trained with four trials for five consecutive days daily. Over the period, mice were conducted to locate and escape onto the platform. The platform position was fixed throughout the test. Each mouse was conducted to swim from a different starting quadrant. Mice failing to swim to the target position within 90 second were guided to the platform and allowed to remain on it for 30 second. In the memory retention session, the hidden platform was removed, the mice were given a time of 60 second to probe, and the time spent in the target quadrant by each mouse was collected.

### Pole test

The pole test for assessing movement disorder was performed as previously described.^20^ A vertical rough-surfaced pole (diameter 1 cm; height 55 cm) was used in this test and the mouse was placed down-upward on its top. The movement time from the pole top to the floor (T-LA time) were measured. The total time was measured with a maximum duration of 30 s.

### Rotarod test

To assess motor coordination in mice, a rotarod test was measured as previously described.^20^ Mice were placed on a rotating bar with a rotation speed of 18 rpm. The time on the rotating bar was recorded as the latent period. Latency to fall was recorded with a maximum of 60 second. Each mouse was given twice consecutive trials and mean value was used to analysis.

### Open field test

Open field test for assessing anxiety-based behaviour was conducted with an open field box (60 × 60 × 40 cm) which was divided in to 9 squares, as previous described.^20^ Each mouse was placed into the centre square. Behaviours were recorded for 6 min. The parameters included total travelled distance and time spent in the central area was used for analysis.

### TUNEL staining and analysis

To evaluate neuronal apoptosis in four different regions, TdT-mediated dUTP-biotin nick end labelling (TUNEL) in conjunction with immunofluorescent staining for NeuN was operated as our previously described.^21^ Briefly, TUNEL staining was performed using an in situ cell death detection kit, following the manufacturer’s protocol. The brain slices were incubated overnight at 4°C with anti-NeuN antibody (1:100) followed by incubation with secondary antibody (1:400) for 2 hours at room temperature. Finally, the slices were stained with DAPI (1:2000). The images were recorded using a confocal microscope (DMi8, Leica, Germany) by an investigator blinded to the experimental design. Similar cerebral regions from six mice in every experimental group were calculated, and the average percentage of TUNEL+/Total DAPI+ was statistically analysed.

### Detection of reactive oxygen species (ROS)

The tissues from frontal cortex (FC), hippocampus (HP), substantia nigra (SN) and striatum (ST) tissues were divided and centrifuged X1000 g at 4°C for 10 minutes. The supernatant was remained and the protein concentration was determined. As the manufacturer’s protocols of ROS assay kit, adding 10 µL of supernatant and 190 µL of BBoxi Probe O12 detector to a 96-well plate, which was incubated for 30 minutes at 37°C in dark. Then fluorescence intensity was measured by fluorescence microplate reader (Infinte M200 Pro, Tecan, Switzerland). The detection result was presented by the ratio of fluorescence value/mg protein.

### Statistical Analysis

We used GraphPad Prism 5 software (GraphPad Software Inc., La Jolla, CA) and SPSS 24 software (International Business Machines Corp., NY, USA) for the data statistical analysis, the level of significance was set at p<0.05. For the case-control study, unpaired two-tailed Student’s t test and *X*^2^ test or Fisher exact test (when expected frequencies of the cells of a contingency table less than 5 were over 20%) were used for analyzing different kinds of data, respectively. For animal experiments, analysis of variance (ANOVA) followed by Fisher’s least significant difference (LSD) or a Tukey-Kramer *post hoc* multiple comparison test for unequal replications was used for statistical analysis.

## ACKNOWLEDGMENTS

This study was supported by Grant No. 81871852 to BL from the National Natural Science Foundation of China, Grant No. XLYC1807137 to BL from LiaoNing Revitalization Talents Program, and Grant No. 20151098 to BL from the Scientific Research Foundation for Overseas Scholars of the Education Ministry of China. grant No. 81200935 to MX from the National Natural Science Foundation of China, Grant No. 20170541030 to MX from the Natural Science Foundation of Liaoning Province. Grant No. 81671862 and No. 81871529 to DG from the National Natural Science Foundation of China.

## AUTHOR CONTRIBUTIONS

M.X, A.V., D.G. and B.L. designed and supervised the study; M.X., S.L., C.D., BN.C., BJ.C., M.J. and W.G. collected the clinic messages and analysed the relevant data; SS.L., Z.L. and M.Z. performed the experiments in vivo and analysed the data; B.L. and A.V. wrote the manuscript.

---

**Supplementary Figure 1.**
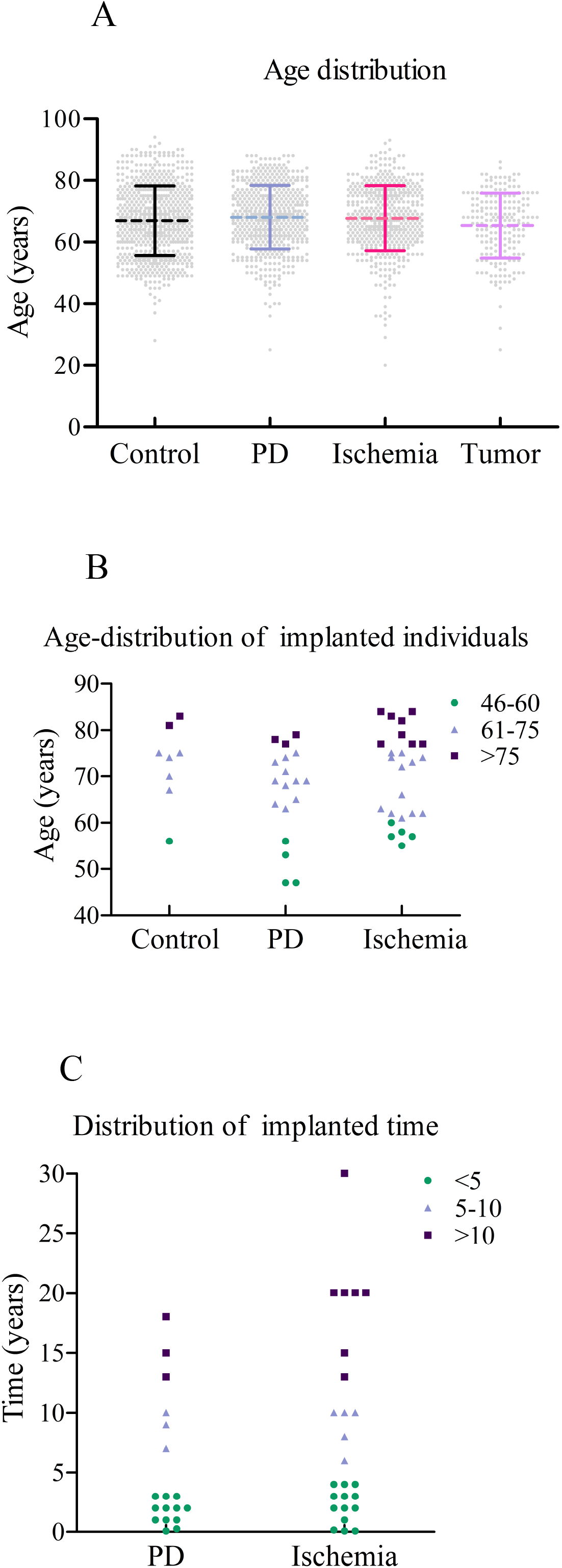
The scatter diagram of implanted age and time in cases of PD and ischemia. **(A)** Scatter diagram of average age in healthy subjects, PD, ischemia and tumour patients. (**B**) The age distribution of the individuals having metal implants. (**C**) The time distribution between metal implanted surgeries and the diagnosed time.

**Supplementary Figure 2.**
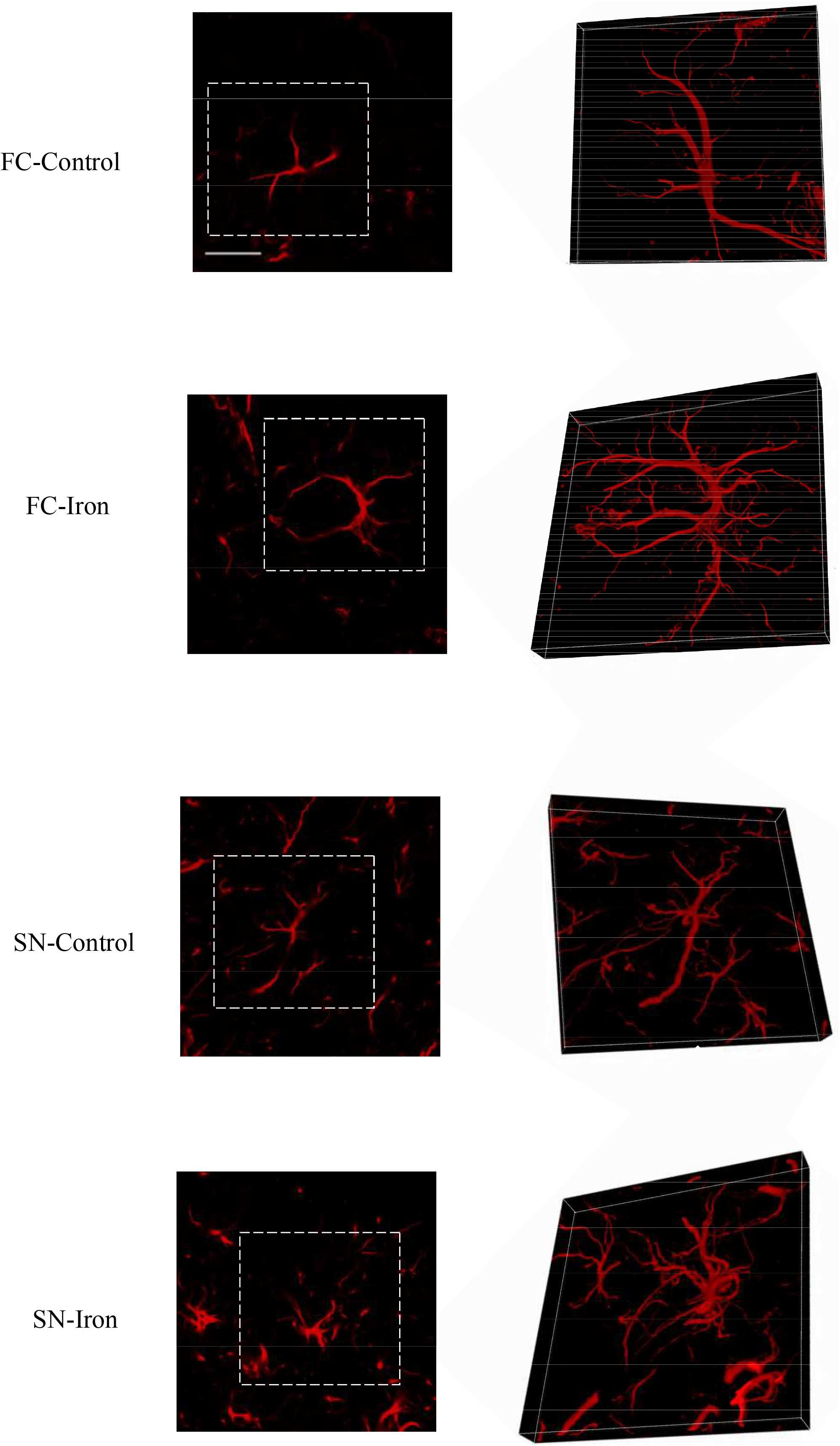
3D-images of GFAP immunofluorescence in frontal cortex and substantia nigra. After treatment with 2 mg/kg/day iron dextran for 6 days, 3D-images of GFAP were taken in FC and SN. Scale bar, 20 μm.

**Supplementary Figure 3.**
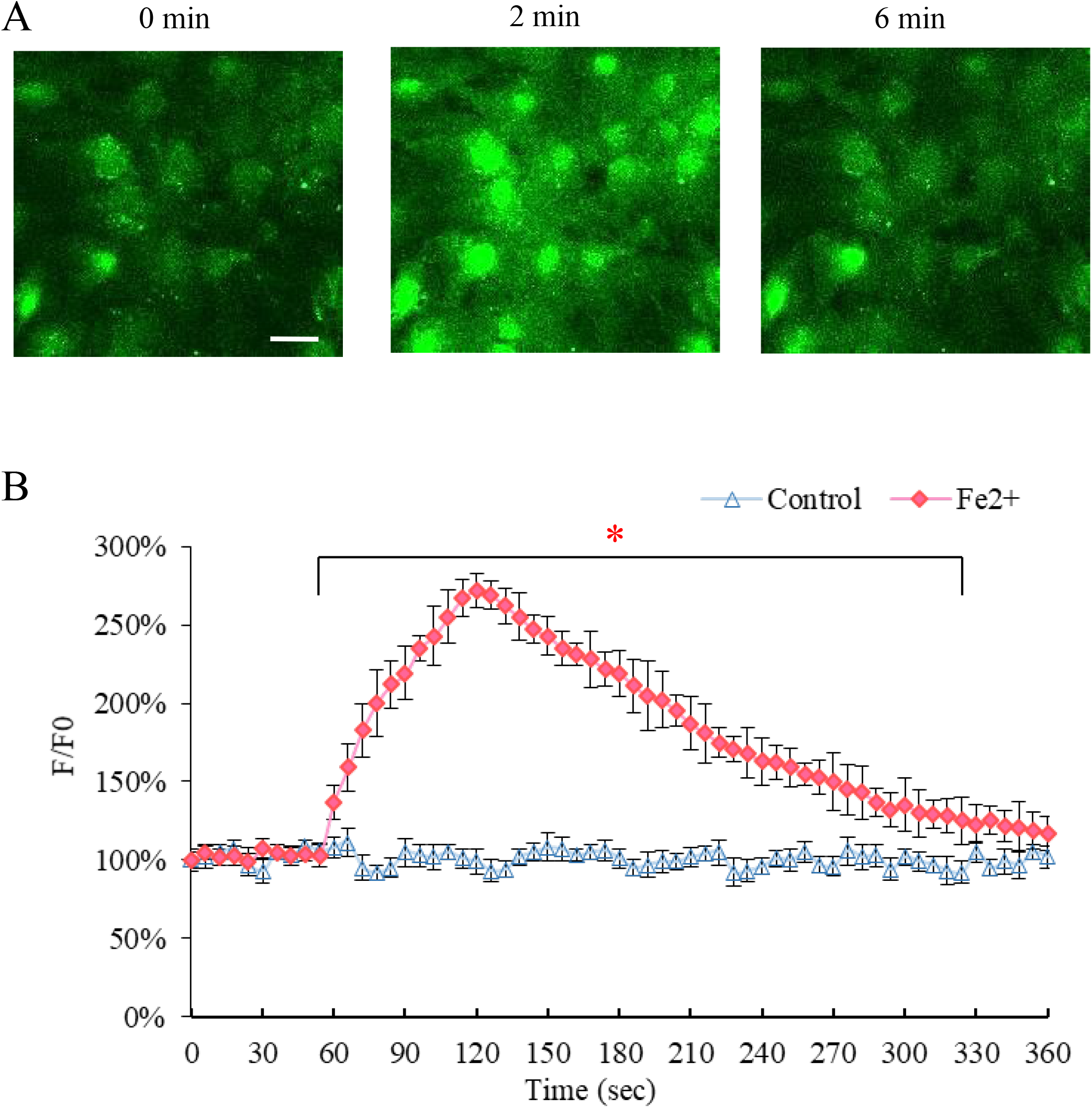
Fe^2+^ directly stimulates intracellular Ca^2+^ signalling. (**A**). Time course of the intracellular Ca^2+^ concentration in response to 1 mM FeSO_4_ in primary cultured astrocytes loaded with fluo-4. Scale bar, 20 μm. The background [Ca^2+^]_i_ was recorded for 1 min, 1 mM FeSO_4_ was added into the medium after 1 min, the representative fluorescence images were shown at 0 min, 2 min and 6 min. (**B**) Traces of fluorescence changes in fluo-4 emission as a function of time after stimulation with FeSO_4_ or saline sodium (control) were measured and represent as mean ± SEM, n = 6.

**Supplementary Figure 4.**
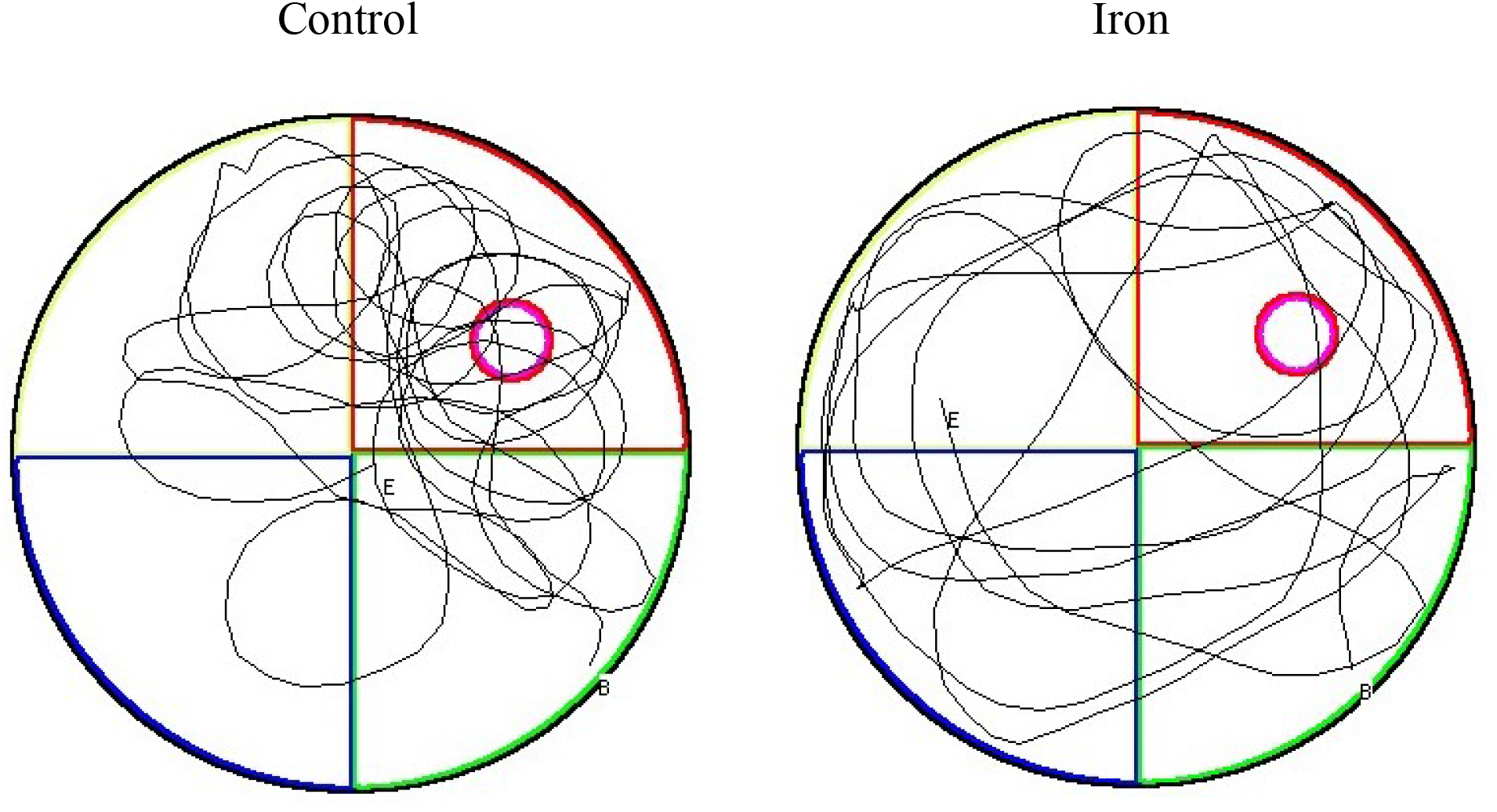
The path diagram of mice in Morris water maze. The representative path diagrams of mice with or without administration of iron dextran.

**Supplementary Table 1.** Metal implants distribution of Parkinson’s disease cases, cerebral ischemia cases and healthy controls.

| Feature | Healthy Control | PD | Cerebra Ischemia |
| --- | --- | --- | --- |
| Total/Inserts number | 700/8 | 500/18 | 500/25 |
| Age at diagnosis (years) |  |  |  |
| <46 | 9/0 | 41/0 | 19/0 |
| 46-60 | 199/1 | 177/4 | 111/5 |
|  |  | P=0.192 | P=0.024 * |
| 61-75 | 322/5 | 244/11 | 259/12 |
|  |  | P=0.036 * | P=0.029 * |
| >75 | 170/2 | 38/3 | 111/8 |
|  |  | P=0.043 * | P=0.016 * |
| Gender |  |  |  |
| Male | 4 (50.00%) | 11 (61.11%) | 13 (52.00%) |
| Female | 4 (50.00%) | 7 (38.89%) | 12 (48.00%) |
|  |  | P=0.683 | P=1.000 |
| Ancestry |  |  |  |
| Han race | 7 (87.50%) | 17 (94.44%) | 25 (100.00%) |
| non-Han race | 1 (12.50%) | 1 (5.56%) | 0 (0.00%) |
|  |  | P=0.529 | P=0.242 |
| Education level |  |  |  |
| Primary and secondary education | 5 | 10 | 16 |
| Skilled vocation and High school education | 2 | 4 | 5 |
| University and postgraduate education | 1 | 4 | 4 |
|  |  | P=1.000 | P=1.000 |
| Time of metal inserts<br>(From surgery to diagnosis) |  |  |  |
| <5 years |  | 12 (66.67%) | 13 (52.00%) |
| 5-10 years |  | 3 (16.67%) | 5 (20.00%) |
| >10 years |  | 3 (16.67%) | 7 (28.00%) |
|  |  | P=0.002 * | P=0.044 * |

